# A minimum threshold for myelination of pyramidal cells in human and mouse neocortex

**DOI:** 10.1101/2023.09.09.556989

**Authors:** M. Pascual-García, M. Unkel, J. A. Slotman, A. Bolleboom, B. Bouwen, A. B. Houtsmuller, C. Dirven, Z. Gao, S. Hijazi, S.A. Kushner

## Abstract

Neocortical pyramidal neurons are frequently myelinated. Diversity in the topography of axonal myelination in the cerebral cortex has been attributed to a combination of electrophysiological activity, axonal morphology, and neuronal-glial interactions. Previously, we showed that axonal segment length and calibre are critical determinants of fast-spiking interneuron myelination (Stedehouder, J. et *al* (2019). However, the factors that determine the myelination of individual axonal segments along neocortical pyramidal neurons remain largely unexplored. Here, we used structured illumination microscopy and cell type-specific manipulations to examine the extent to which axonal morphology determines the topography of axonal myelination in mouse neocortical pyramidal neurons. We found that, unlike what was determined for fast-spiking interneurons, the joint combination of axonal calibre and interbranch distance does not predict axonal myelination in pyramidal neurons, rather it provides a minimum threshold for myelination; pyramidal neurons with an axon calibre and interbranch distance lower than 0.24 µm and 19 µm, respectively, are almost never myelinated. Moreover, we further confirmed that these findings in mice also extend to human neocortical pyramidal cell myelination, suggesting that this mechanism is evolutionarily conserved. Taken together, our findings suggest that axonal morphology is highly deterministic of the topography and cell-type specificity of neocortical myelination.

## Introduction

Diverse neuronal types are synaptically connected within the cerebral cortex forming complex and interactive circuits between different subcortical layers and brain regions. Axonal myelination is a crucial mammalian neurobiological adaptation that functions as an electrical insulator, enabling saltatory conduction of action potentials^1^, facilitating reliably timed neuronal activity^2,3^, and optimizing neuronal energy expenditure^4^. Accordingly, loss or damage to the myelin sheath or oligodendrocyte integrity has been shown to contribute to a variety of neuropsychiatric disorders^5^.

Myelin composition varies across brain regions. Developmentally, CNS myelination exhibits protracted maturation throughout childhood, adolescence and early adulthood, with considerable cell-type heterogeneity, as well as across distinct subcortical layers and brain areas^6,7^. In particular, the topography of myelination along individual axons of neocortical pyramidal neurons is known to be highly heterogeneous^8 9,10^. This has raised the question of how oligodendrocytes determine their targets among the totality of axons. Intrinsic molecular cues from axons have been suggested to inhibit or attract myelinating oligodendrocytes^11,12^. Moreover, neuronal activity has been shown to contribute to the development of oligodendrocytes and therefore the sheathing of axons^13^. However, although neocortical pyramidal cell axons are a well-characterized target of myelinating oligodendrocytes, the heterogeneity of their internodal topography remains poorly understood.

Axonal diameter is widely known to be an important determinant of myelination. Schwann cells, the myelinating cells in the peripheral nervous system, almost exclusively ensheath axons with a diameter greater than ∼1 µm^15–17^. In the CNS, oligodendrocytes restrict their ensheathment to axons with a diameter greater than ∼0.3 µm^10,18,19^. However, many axons exceeding this minimum threshold remain unmyelinated^19^. This suggests that in the CNS, other factors influence axonal myelination. Previous studies of local fast-spiking interneurons in the cerebral cortex demonstrated that axonal morphology, including calibre and interbranch segment length, is sufficient to predict interneuron myelination with high accuracy^19^. Whether pyramidal cells adhere to similar or distinct rules governing their myelination has yet to be determined.

Here, we investigated the relationship between pyramidal cell axonal morphology and myelination in layer II/III of mouse somatosensory and prefrontal cortices. We observed that unlike fast-spiking interneurons, the joint combination of diameter and axonal segment length is not predictive of segmental myelination in the somatosensory cortex (S1) but rather imposes a necessary threshold for myelination, deterring the myelination of axons with a calibre and length that are lower than the identified respective thresholds . We found that, in the prefrontal cortex (PFC), pyramidal cells are rarely myelinated at 8-12 weeks of age, likely dure to their immature axonal morphology. Lastly, using human *ex vivo* neurosurgically resected tissue, we confirmed that the morphological thresholds for neocortical pyramidal cell myelination in mice also extend to human. Taken together, local axonal morphology of pyramidal cells appears to explain a substantial proportion of the variance underlying segmental myelination and might function as a causal biophysical constraint for oligodendrocytes to initiate axonal myelination.

## Results

### Myelination of neocortical pyramidal cells in the somatosensory cortex is associated with axonal morphology

To establish the relationship between axonal morphology and myelination of pyramidal cells in the somatosensory cortex (S1), we performed whole-cell patch-clamp recordings in S1 slices from wild-type (WT) mice at 8-12 weeks (**Figure 1a, b; Supplementary table 1a**). Biocytin-labelled neurons were imaged using confocal microscopy and immunostained with myelin basic protein (MBP) to investigate their myelination profile (**Figure 1c**). 80% (8 out of 10) of the examined cells were myelinated. Consistent with previous studies^9^, only the primary axon had myelinated segments (**Figure 1d; Supplementary table 1b**). Axonal diameter and length of each reconstructed segment were measured using SIM imaging. The axonal shaft diameter, independent of myelination status, had an average diameter of 0.262 µm (**Figure 1e**). Importantly, segments that exhibited myelination had a larger calibre compared to those that were unmyelinated. Moreover, myelinated segments were longer on average than those that were unmyelinated (**Figure 1**f, g).

**Figure 1.**
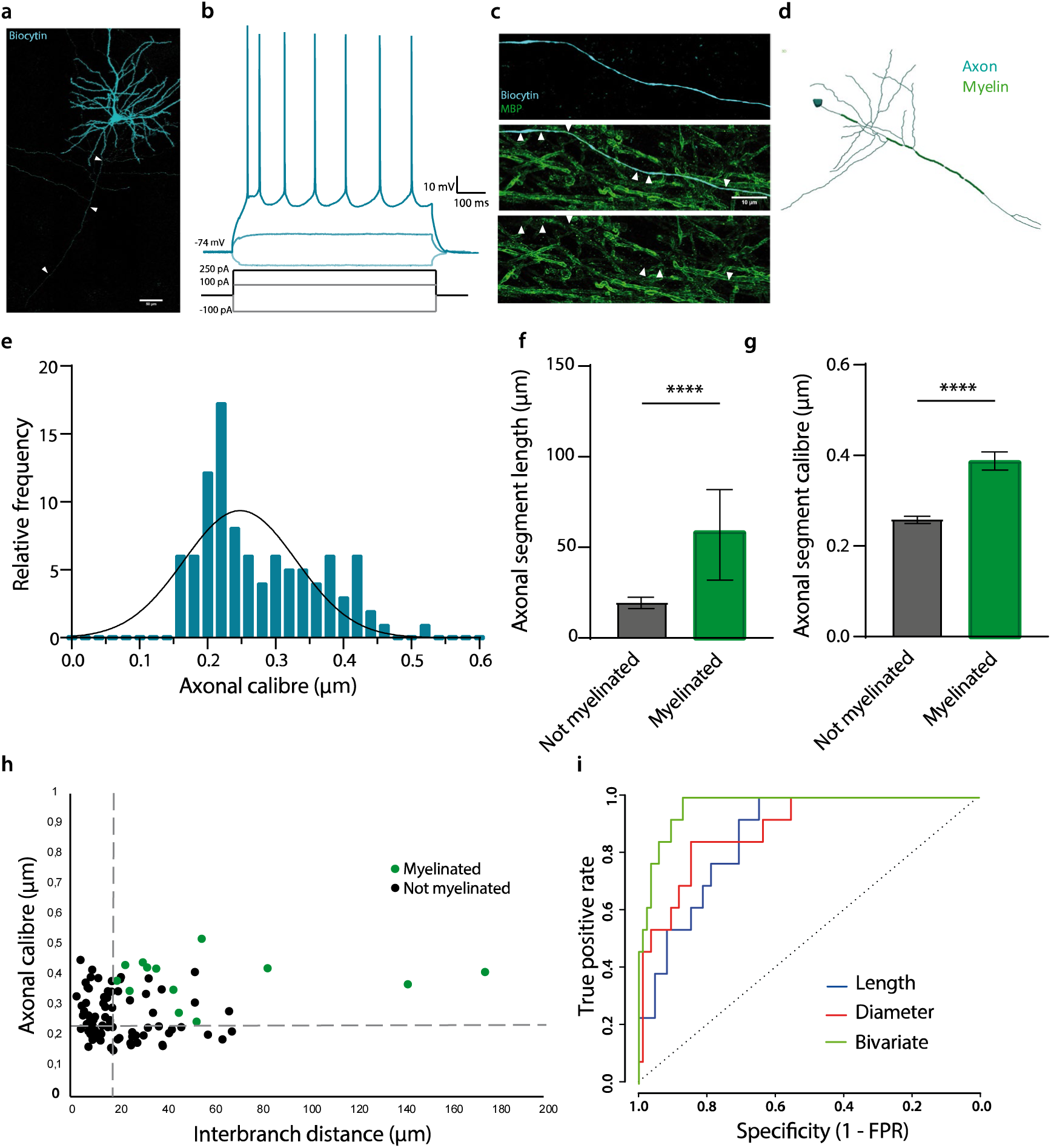
Axonal properties of pyramidal cells in the mouse S1. **a)** Example of a pyramidal cell in S1 filled with biocytin. **b)** Electrophysiological example traces of a layer II-III pyramidal cell at hyperpolarising (light blue) and depolarising currents (blue and dark blue). **c)** High magnification picture of a myelinated segment (biocytin in blue; MBP in green). Arrow heads indicate myelinated segments. **d)** Neurolucida reconstruction of a representative S1 pyramidal cell including the myelinated segments in the main axon (axon in blue; myelin in green). **e)** Frequency histogram of the axonal segment calibre, fitted with a Gaussian curve; mean = 0.28 µm ± 0.008 µm s.e.m.; n = 98 axonal segments from 10 cells. **f)** Comparison of the axonal segment length between myelinated and unmyelinated segments (Mann-Whitney: U = 149; p-value < 0.0001; unmyelinated segments = 14.71 µm; myelinated segments = 43.56 µm; n = 98 segments). **g)** Calibre of myelinated and unmyelinated segments (t-test: t = 5.851; p-value < 0.0001; unmyelinated segments = 0.26 µm; myelinated segments = 0.39 µm; n = 98 segments). **h)** Distribution of the axonal segment calibre and interbranch distances for myelinated (green) and unmyelinated (black) segments. Dotted lines depict the bivariate thresholds indicated in figure **i)** ROC curves for univariate (length or calibre) and bivariate analyses.

Using receiver operating characteristic (ROC) analysis, we examined the sufficiency of axonal calibre and length to predict segmental myelination in S1 pyramidal neurons. As expected, axonal diameter was a significant predictor of segmental myelination with a critical threshold at 0.346 µm (AUC = 0.97; sensitivity = 0.85; specificity = 0.85). Axonal length was also significantly associated with segmental myelination, albeit with a somewhat lower specificity (threshold, 18.90 µm; AUC = 0.87; sensitivity = 1; specificity = 0.65). Therefore, we performed a bivariate ROC analysis as previously described^19^, in order to determine whether the joint combination of axonal calibre and length might yield better estimates of segmental myelination along pyramidal cell axons. The combination of both parameters yielded a significant yet mild improvement in the prediction of segmental myelination (AUC = 0.97; sensitivity = 1; specificity = 0.88), with thresholds for axonal calibre and length of 0.236 µm and 18.14 µm, respectively (**Figure 1h, i**; axonal calibre: p < 0.01; axonal length: p < 0.01). Importantly, the specificity and accuracy of the prediction was still below 90%. This is demonstrated by the many unmyelinated segments that crossed the identified threshold (10/24). Given that none of the segments below the threshold were myelinated, this data suggests that our bivariate model presents a threshold that is necessary for myelination but not critical; pyramidal neurons with a joint combination of axonal calibre and length that are below 0.24 µm and 19 µm respectively will not be myelinated.

### Prefrontal cortex layer II-III pyramidal cells are seldomly myelinated in mice

Next, we aimed to investigate whether the above observations were similar in a different neocortical region. Using the same workflow as in S1, we patched and filled pyramidal cells with biocytin in the prefrontal cortex (PFC) of naïve wild-type mice (**Figure 2a, b; Supplementary table 2**). The targeted cells were immunostained with MBP to study their myelination profile. 95% (17 out of 18) of the cells showed no myelination, and 1 cell showed one myelinated segment on the totality of its axon (**Figure 2c-e**). This is consistent with earlier reports that myelination in the PFC occurs later in development than other neocortical regions^20^. Does this lack of myelination in PFC pyramidal neurons reflect an immature morphological profile of their axons compared to S1? To answer this, we performed structural illumination microscopy (SIM) and measured the diameter of individual axonal segments of the reconstructed PFC cells. We found that the distribution of axonal calibre was significantly different in S1 when compared to PFC; only 4.1% of PFC segments had a calibre larger than 0.4 µm whereas 12.8% of S1 segments were above 0.4 µm (**Figure 2f,g**). It is also intresting to note that the biggest shift in the curve was above the minimum threshold of 0.24 µm (dashed green line, **Figure 2g**). No significant difference was observed in the distribution of the axonal segment’s length, suggesting that these two variable alone can’t explain the difference reported in the percentage of myelinated pyramidal cells in S1 versus PFC. Unsurprisingly, the necessary threshold obtained for S1 (axonal calibre below 0.24 µm and length below 19 µm) was applicable for PFC pyramidal neurons as none of the segments examined were myelinated.

**Figure 2.**
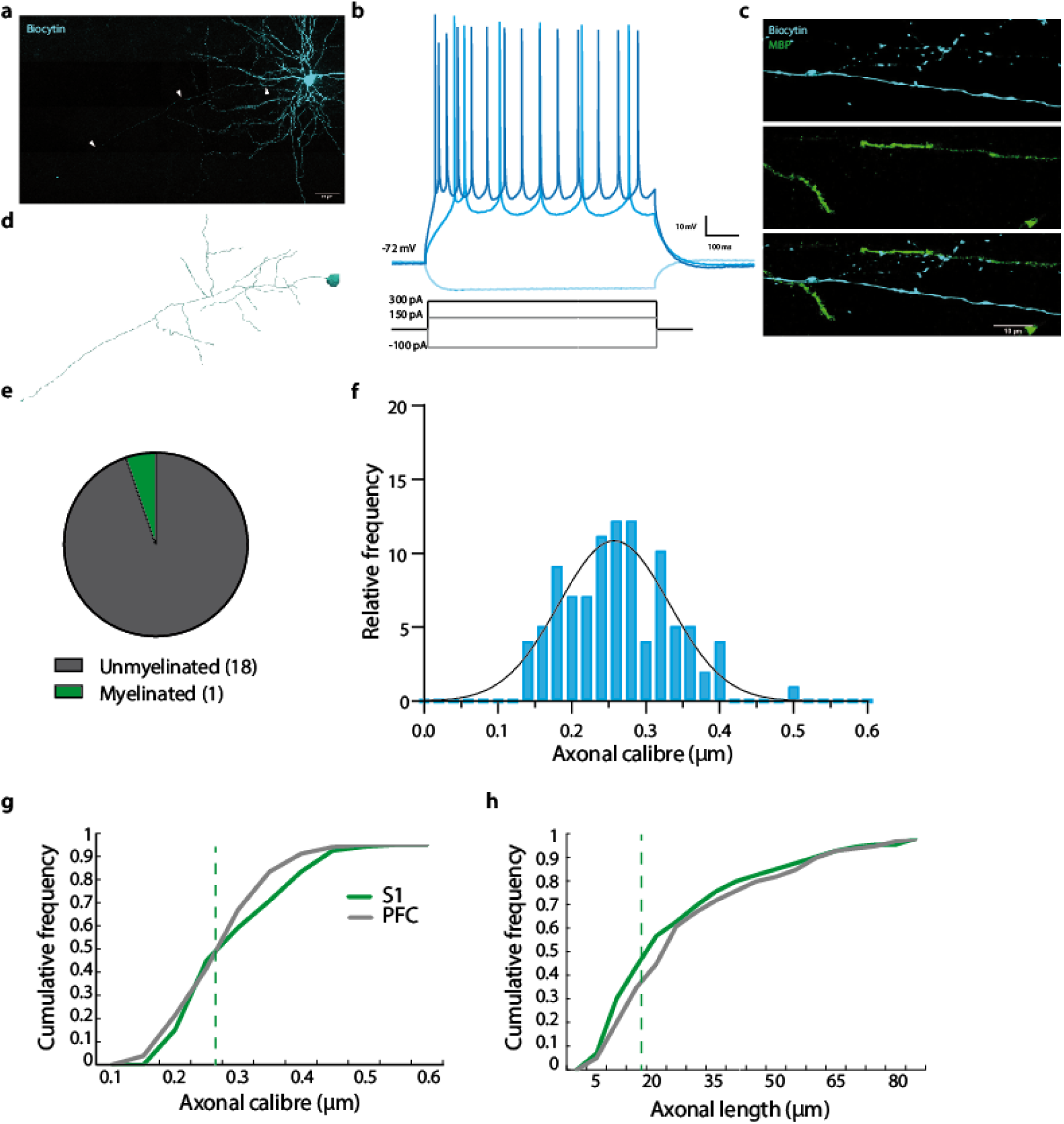
Axonal properties of pyramidal cells in the PFC. **a)** Pyramidal cell filled with biocytin (blue) and immunostained with MBP (green) to visualise its myelination. Arrow head indicates main axon. **b)** Electrophysiological example traces of a layer II-III pyramidal cell at hyperpolarising (light blue) and depolarising currents (blue (rheobase) and dark blue). **c)** High resolution picture of an unmyelinated segment of a pyramidal cell in PFC (cell in blue and MBP in green). **d)** Neurolucida reconstruction of the pyramidal cell in a). **e)** Proportion of myelinated cells in the PFC. **f)** Histogram of the distribution of axonal segment calibre fitted with a Gaussian curve; mean = 0.26 µm ± 0.0073 µm s.e.m; n = 98 axonal segments from 19 cells. **g)** Cumulative frequency distribution of axon calibre of all segments from S1 (green) and PFC (grey). **h)** Cumulative frequency distribution of axon length of all segments from S1 (green) and PFC (grey). The dashed line indicates the minimum myelination threshold.

### Genetic enlargement of PFC pyramidal cell size increases their likelihood of myelination

Prior studies have demonstrated that increasing cell size can promote myelination of the individually targeted cells^19,21^. Given that we observed nearly no myelinated segments in PFC pyramidal cells at 8-12 weeks, we wondered whether increasing their size might alter the minimum threshold and facilitate their myelination. To examine this possibility, we induced a cell-type specific deletion of the *Tsc1* gene, a negative regulator of the mTOR signalling pathway and determinant of cell size^22^ (**Supplementary figure 1a**). At P28, adeno-associated virus (AAV9) expressing cre recombinase with an N-terminal GFP tag (GFP-Cre) under the control of the αCaMKII promotor was stereotactically injected into the PFC of one hemisphere of homozygous floxed *TSC1^f/f^* mice, while the contralateral PFC was injected with a control virus that expressed GFP alone (**Figure 3a**). In both hemispheres, all GFP expressing cells had a pyramidal morphology, consistent with the cell-type selectivity of the αCaMKII promotor^23^ (**Figure 3b, 4c**). Moreover, cells in the hemisphere expressing GFP-Cre exhibited elevated pS6, consistent with deletion of the *Tsc1* gene, in contrast to the contralateral hemispheres expressing GFP alone (**Supplementary figure 1b**). Accordingly, as expected, GFP-Cre expressing pyramidal cells, but not those expressing GFP alone, exhibited a pronounced increase in soma area (**Figure 3b, c**).

**Figure 3.**
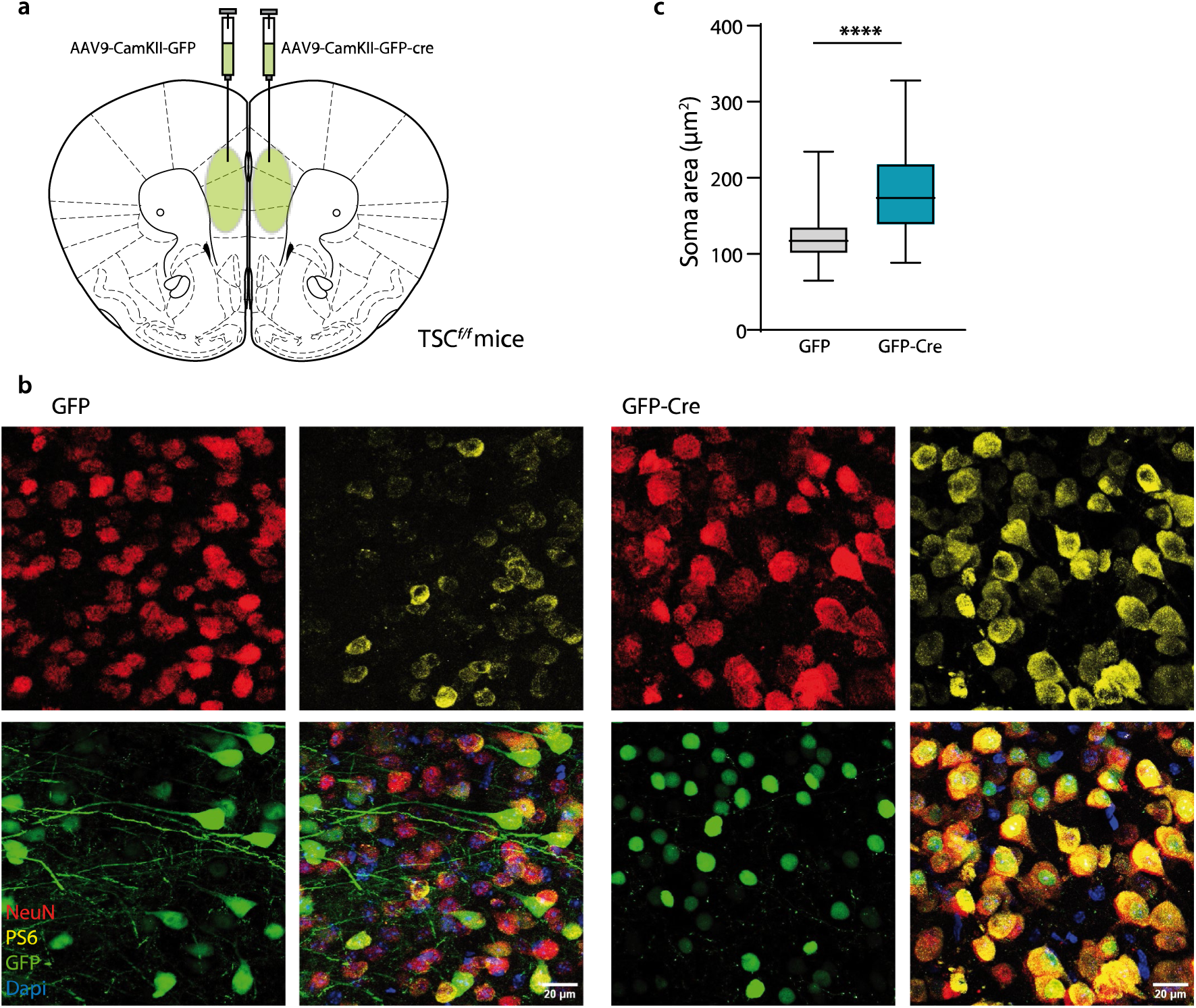
*Tsc1* deletion enlarges pyramidal cells. **a)** 4-weeks TSC1*^f/f^* mice were virally injected with unilateral CamKII-GFP-cre and contralateral CamKII-GFP viruses in the PFC. One month later mice were sacrificed are analysis were carried out. **b)** Confocal images of the area of injection at 40x resolution depicting colocalization of NeuN (red), PS6 (yellow), GFP (green) and Dapi (blue). GFP in GFP-Cre cells is only visible in the nucleus because Cre expression is nuclear, whereas GFP in GFP cells is found along the whole cytoplasm. Scale bar 20 µm. **c)** Box plot of the area of the soma of NeuN- and GFP-positive cells (Mann-Whitney: U = 5625; p-value < 0.0001; GFP average: 117.2 µm^2^, n = 163 cells from 4 mice; GFP-Cre average: 173.5 µm^2^, n = 207 cells from 4 mice).

Next, we investigated whether pyramidal cell-specific deletion of TSC1 enhanced their myelination. Therefore, we patched and filled GFP-labelled cells in the PFC, and stained for MBP (**Figure 4a-c**). GFP-Cre neurons showed decreased excitability; the action potential (AP) threshold was increased and the firing frequency was reduced at small depolarising steps, compared to GFP neurons (**Supplementary figure 2a**). Moreover, along with their increased size, both the capacitance and axonal calibre of GFP-Cre cells were significantly increased compared to GFP control cells (**Figure 4d; Supplementary figure 2a, b**). Consistent with the increased axonal calibre, the proportion of myelinated GFP-Cre cells increased substantially compared to GFP cells (GFP-Cre, 10 of 16 cells, 62.5%; GFP alone, 5 of 19 cells, 26.3%; *P* < 0.05, Fisher’s Exact Test; **Figure 4e**). Importantly, the cut-off threshold for myelination held true for both data sets; only 1 segment out of 30 in GFP-Cre and 1 out of 40 in GFP mice with an axon calibre below 0.24 µm and a length less than 19 µm were myelinated. The increased number of myelinated segments in the GFP-Cre mice was driven by the increased calibre size as most of the myelinated segments had an axonal diameter larger than 0.45 µm.

**Figure 4.**
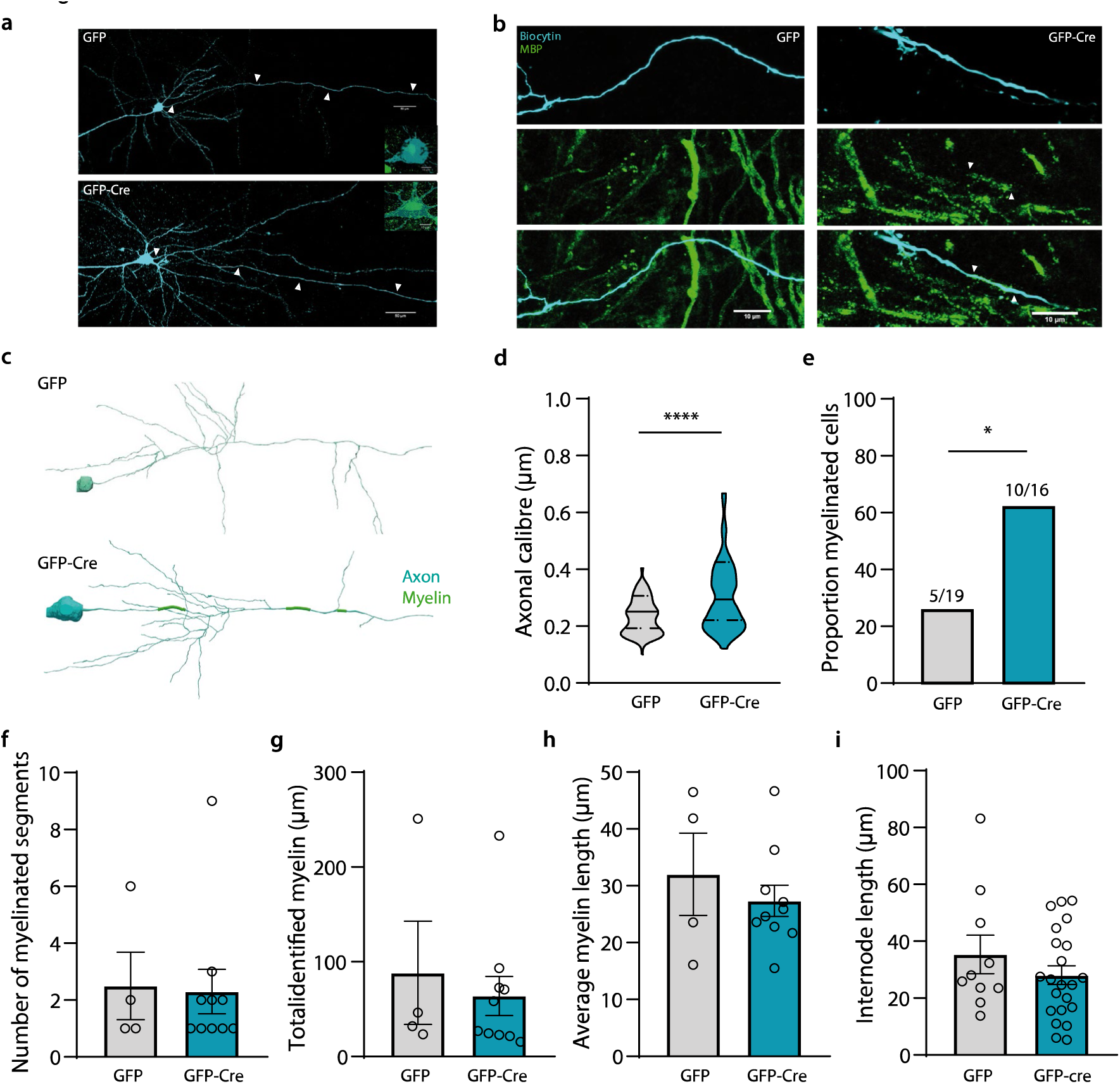
Axonal properties of pyramidal cells in the PFC. **a)** Representative example of filled PFC pyramidal cells. Arrow heads indicate the main axon. Zoom in of colocalization of GFP with biocytin-filled cells. **b)** High magnification confocal image of biocytin-labelled GFP (left) and GFP-Cre (right) pyramidal cells (dark blue) and its myelinated segments (MBP in green). Arrow heads indicate the myelinated segments. Scale bar 10 µm. **c)** Example of Neurolucida axonal reconstruction of an unmyelinated GFP pyramidal cell (top) and a myelinated GFP-Cre pyramidal cell (down). **d)** Violin plot showing that axonal calibre is larger in GFP-Cre cells compared to GFP cells (Mann-Whitney: U = 4296; p-value < 0.0001; GFP average: 0.25 µm, n = 104 segments from 9 cells; GFP-Cre average: 0.30 µm, n = 120 segments from 13 cells). **e)** The proportion of myelinated cells is increased in GFP-Cre cells compared to GFP cells (Fisher’s exact test: p-value = 0.044; GFP-Cre: 62.5% (10 out of 16); GFP: 26.3% (5 out of 19)). **f)** Number of myelinated segments is the same in both groups (t-test: t = 0.1389; p-value = 0.89; GFP average: 2.5 myelinated internodes; GFP-Cre average: 2.3 myelinated segments). **g)** Total amount of myelin is the same across groups (t-test: t = 0.5245; p-value = 0.61; GFP average: 88.37 µm; GFP-Cre average: 64.01 µm). **h)** Average myelin length is similar between groups (t-test: t = 0.7572; p-value = 0.46; GFP average: 32.01 µm; GFP-Cre average: 27.35 µm). **i)** The average internode length remains the same in both conditions (t-test: t = 1.094; p-value = 0.29; GFP average: 35.35 µm; GFP-Cre average: 28.06 µm). nGFP = 5 cells; nGFP-Cre = 10 cells.

To determine whether the increase in the proportion of myelinated GFP-Cre cells also induced changes in the topography of myelination along individual axons, we performed Neurolucida reconstructions of filled cells and quantified the number and length of internodes (**Figure 4c**). We found no differences in the number of myelinated segments, the total amount of myelin per cell or the average internode length (**Figure 4f-i**). Together, these data show that enlargement of pyramidal neurons increases the proportion of myelinated axons due to an increase in their axonal calibre, but without an overall increase in myelination per cell.

### Human neocortical pyramidal neurons exhibit a similar morphological relationship with axonal myelination

As our previous data uncovered a predicative role for axonal morphology in the myelination of fast-spiking interneurons in mice and in humans, we next wondered if what the minimum threshold observed in mice pyramidal neurons holds true for human pyramidal neurons. To that end, we performed whole-cell electrophysiology and biocytin-filling of patched human pyramidal cells in neurosurgically resected neocortical tissue (9 cells from 5 patients; see *Methods*). Pyramidal cells were visually targeted by their stereotypical morphology. All recorded cells exhibited canonical electrophysiological characteristics of pyramidal neurons such as regular firing frequency, high amplitude and the subthreshold depolarisation (**Figure 6a, b; Supplementary table 3a**). *Post hoc* MBP immunofluorescence revealed that 89% of the cells (8 out of 9) were myelinated with an average internode length of 49.37 ± 12.69 µm, total myelin length of 162.21 ± 82.3 µm and 2.87 ± 1.25 number of internodes (**Figure 6c, d; Supplementary table 3b**).

**Figure 6.**
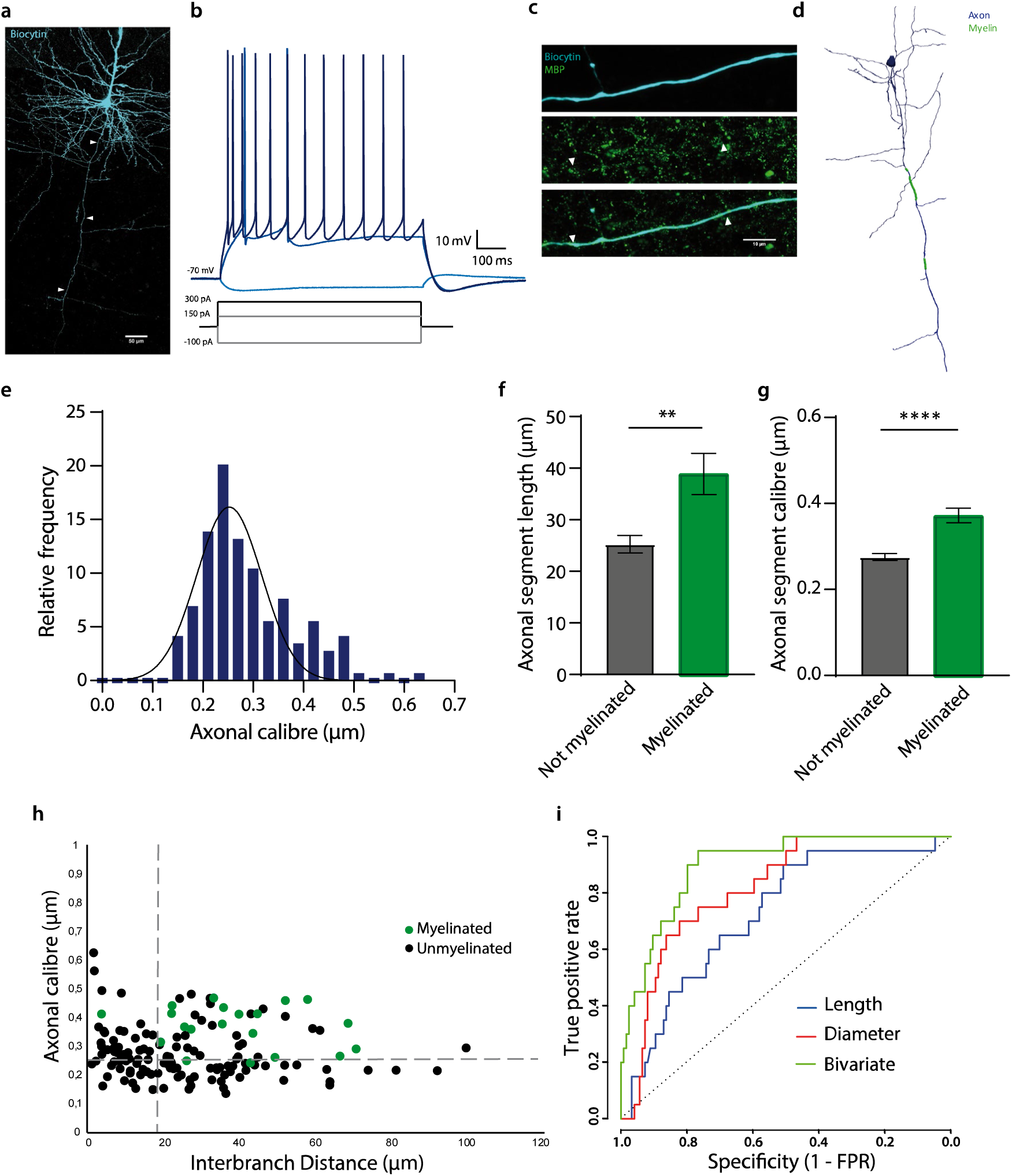
Axonal properties of human pyramidal cells. **a)** Confocal image of a representative human pyramidal cell filled with biocytin. **b)** Electrophysiological example traces of a human pyramidal cell at different hyperpolarising (light blue) and depolarising currents (blue (rheobase) and dark blue). **c)** High resolution image of a myelinated segment (MBP; green) in a human pyramidal neuron (dark blue). Arrowheads delimit myelin internode. **d)** Neurolucida axonal reconstruction of a representative pyramidal cell (axon in dark blue; myelin in green). **e)** Frequency histogram of the axonal segment calibre, fitted with a Gaussian curve; mean = 0.29 µm ± 0.008 µm s.e.m.; n = 144 axonal segments from 10 cells. **f)** Axonal segments that were myelinated were significantly longer than those that did not present any internode ((t-test: t = 2.989; p-value = 0.003; unmyelinated segments: 25.28 µm; myelinated segments: 38.88 µm). **g)** Axonal calibre of myelinated segments is larger than those unmyelinated (t-test: t = 4.602; p-value < 0.001; (unmyelinated segments: 0.27 µm; myelinated segments: 0.37 µm). **h)** Distribution of the axonal segment calibre and interbranch distances for myelinated (green) and unmyelinated (black) segments. Dotted lines depict the critical thresholds of the bivariate analysis. **i)** ROC curves for univariate (length and calibre) and bivariate analyses.

Super-resolution analysis of individual axonal segments yielded a mean axonal shaft diameter of 0.289 ± 0.77 µm (**Figure 6e**). Similar to our observations in S1 pyramidal cells in mice, human neocortical pyramidal cells revealed a strong association between segmental myelination and the joint combination of axonal diameter and axonal length. Segments that lacked myelination exhibited shorter length and thinner diameters compared to those that were myelinated (**Figure 6f, g**). ROC analysis of the univariate models revealed thresholds of 21.75 µm for segment length (AUC = 0.73; sensitivity = 0.90; specificity = 0.51) and 0.348 µm for axonal diameter (AUC = 0.82; sensitivity = 0.70; specificity = 0.82, respectively). The bivariate analysis that included the joint combination of axonal calibre and segment length revealed combined thresholds of 18.96 µm for segment length and 0.246 µm for axonal calibre (AUC = 0.90; sensitivity = 0.95; specificity = 0.77). The joint combination of both parameters significantly improved the ROC model compared to univariate models based on axonal calibre (*P* = 0.027) or segment length (*P* < 0.001) (**Figure 6h, i**). Almost all the myelinated segments (1 out of 21) were above the necessary threshold for myelination. It is interesting to note that S1 pyramidal neurons from mice and human pyramidal neurons adhere to a very similar limiting threshold (calibre 0.24 µm in mice vs 0.25 µm in humans; length 18.14 µm in mice vs 18.96 µm humans), and show a comparable percentage of myelinated pyramidal cells. Together, these data indicate that myelination adheres to similar morphological thresholds across different species.

## Discussion

Pyramidal cells are the major class of excitatory glutamatergic neurons in the cerebral cortex. Their function, morphology, excitability and number vary across different layers of the neocortex. The myelination profile of pyramidal cells is highly diverse and depends on their electrical activity and connectivity^9^. However, few studies have previously been performed to investigate the cellular determinants underlying the spatial distribution of neocortical pyramidal cell internodes. In the present study, we provide new insights regarding the relationship between axonal morphology and the myelination of individual segments along pyramidal cell axons. We employed cell-specific techniques to assess the causality of this relationship by genetically targeting individual pyramidal cells and using high-resolution methods to measure axonal diameter and length. Our findings reveal a cut-off threshold for the myelination of pyramidal cell axonal segments lower than 18.5 µm and thicker than 0.24 µm. These results also provide an explanation as to why only the primary axon is myelinated in neocortical pyramidal cells, as collateral branches protruding from the main axon of pyramidal neurons have a notably thinner diameter, which is typically subthreshold for myelination of these segments. Notably, the lack of myelination along branches of pyramidal cells is in marked contrast to collateral branches of fast-spiking interneurons, which are frequently myelinated in both mouse and human neocortex^19,24^.

### Differential myelination profile across brain regions

Internodal topography varies by neuronal type, cortical layer and brain region^25^. Cortical layers V-VI, for instance, are densely myelinated in comparison to more superficial layers^9^. Consistent with earlier studies, we found that pyramidal cells in layer II-III of S1 were frequently myelinated. Myelinated segments of these cells were on average longer and thicker than those that were unmyelinated (**Figure 1**).

In contrast to the findings in S1, we found that pyramidal cells in layer II-III of the PFC were rarely myelinated in naïve mice at 8-12 weeks of age. Interestingly, myelin sheaths were detectable on other neuronal types, such as local interneurons in the same area^24^, indicating that the lack of myelin on pyramidal neurons is not due to an absence of myelinating oligodendrocytes in this region. Why are pyramidal neurons not myelinated in the PFC at this age? One explanation could be that these cells are not yet fully mature. The PFC is critically involved in complex cognitive and executive function^27^, and developmentally one of the last regions to mature, undergoing continual changes through adolescence until early adulthood^28^. It is well-established that the PFC in humans is one of the last regions to become fully myelinated^29,30^. Moreover, cortical myelination in mice exhibits a similar developmental trajectory, peaking at one year of age^31,32^. Electrical cues acquire critical relevance during development, and activity-dependent myelination and crucial for remodelling the brain’s networks^14^. Another likely explanation is that, as the mice used in the study were naïve young adults, very little experience-dependent myelination had occurred in the PFC at this stage. It is interesting to note that, in mice injected with the control GFP virus, 5 out of 19 pyramidal cells showed myelination, as opposed to what was observed in naïve untreated mice (1 out of 18), suggesting a role for intracranial injections in the myelination of axons. In fact, mice that received sham intracranial injection (no virus injected) also showed some myelinated segments in PFC pyramidal neurons (data not shown).

### Cellular morphology is linked to excitability and myelination

Our previous work demonstrated that enlargement of neocortical fast-spiking interneurons by deletion of *Tsc1* increased the number and length of internodes in these cells^19^. Likewise, manipulation of pyramidal cells in *Tsc1^f/f^* mice increased cell size, including axonal calibre. The proportion of GFP-Cre pyramidal cells that were myelinated increased substantially in PFC. These data suggest that *de novo* myelination may be induced in unmyelinated cells by altering their axonal morphology but does not affect the number of pre-existing internodes. Notably, the required distance between sodium clusters in paranodal regions is on average ∼32 µm, which might explain why the minimum segmental length is at least 17 µm^33^ for oligodendrocytes to ensheath axons.

Neuronal excitability is another factor to take into consideration as a potential determinant of myelination profiles. Adaptations of axonal diameter have been observed in response to neuronal stimulation^34^. Neuronal activity enhances myelin formation and regulates the complexity of axonal arborization as well as the thickness and number of myelinated segments in both excitatory and inhibitory neurons^14,35,36^. Previous studies have observed a relationship between pyramidal cells’ firing frequency and larger axonal diameter, which serves to reduce their energetic consumption by enhancing their axonal myelination^26,37^. Notably, we observed that enlarged GFP-Cre pyramidal cells in the PFC exhibited reduced excitability, with a higher AP threshold and reduced peak and firing properties compared to their control littermates (**Supplementary figure 2a**). Together, we find that, at the individual cell level, the relationship between axonal myelination and experimental manipulation of pyramidal cell size more closely follows changes in morphology rather than excitability, indicating that myelination is not only driven by activity but also by morphological plasticity.

### Biophysical cues for oligodendrocyte attraction

Extrinsic and intrinsic cues guide oligodendrocyte precursor cells (OPCs) to mature into myelinating oligodendrocytes and ensheath axons^38^. Beyond fibre diameter, axon curvature has also been suggested to play an important permissive role in myelination^39^. Bechler, *et al*. demonstrated that oligodendrocytes increase the sheath length in larger fibres^40^. Our results indicate that axonal length also creates a physical constraint for myelination, given that segments shorter than 18 µm are mostly unmyelinated. Furthermore, axonal branch points are consistently unmyelinated, which might suggest that the angle formed by the emerging daughter branches biophysically restricts oligodendrocyte ensheathment. Consistent with this finding, branch points of myelinated axons are also common locations for nodes of Ranvier and therefore critical for signal integration along the axon^41,42^.

### Pyramidal cell myelination is stable across species

Pyramidal cells in the human neocortex are extensively myelinated^9^. Single-cell axonal reconstruction of pyramidal cells in different brain areas, including frontal and temporal cortices, showed that almost 90% of human pyramidal cells were myelinated. This increased myelination ratio in humans when compared to mice could be due to a mere species difference or another more likely explanation is that it is the result of a difference in learning-induced myelination in humans, as all the mice used in this experiment were naïve young adults whereas the mean age of the patients in this study was 55 years old. Interestingly, bivariate analysis revealed human myelination thresholds of 18.96 µm for segment length and 0.25 µm for axonal diameter, which are very similar to what we found in S1 cells from mice (18.14 µm and 0.24 µm, respectively). Such similarities suggest that oligodendrocytes have intrinsic mechanisms for myelination that are highly preserved across species and guarantee the correct functioning of the network. Likewise, fast-spiking interneurons also exhibit highly similar morphological thresholds^19^, suggesting that oligodendrocyte ensheathment potential might be independent of neuronal cell type and strongly determined by biophysical constraints.

In conclusion, pyramidal cell axonal morphology is a major determinant of myelin topography. Moreover, the predictive thresholds for axonal calibre and length are similar across multiple regions in mouse and human neocortex, suggesting the possibility of an evolutionarily conserved biophysical mechanism governing CNS axonal myelination.

## Materials and Methods

### Mice

All experiments were conducted under the approval of the Dutch Ethical Committee and in accordance with the Institutional Animal Care and Use Committee (IACUC) guidelines.

For this study, two different mouse lines, obtained from Jackson Laboratory, were utilised: *C57BL/6J* (referred as WT; strain #000664) and *Tsc1^tm1Djk^/J* (referred as *Tsc1^f^*^/*f*^; strain #005680). All lines were backcrossed for more than 10 generations in C57BL/6J. Inducible TSC1 knockout mice were generated by crossing conditional biallelic floxed *Tsc1* mutant heterozygous mice (*Tsc^f/f^*) in C57BL/6J background. Mutant lines were backcrossed to obtain homozygous *Tsc^f/f^* mice. All mutant lines were viable and healthy. No behavioural abnormalities and no spontaneous seizures were ever observed.

Mice between 8 and 12 weeks old were used in these experiments. All groups consisted of mice from both sexes. WT animals were group-housed whereas those animals that underwent surgical procedure were single-housed after the viral injection. All mice were maintained on a regular 12 h light/dark cycle at 22 °C (±2 °C) with access to food and water *ad libitum*.

### Human

Peri-tumoral neocortical tissue was obtained from 5 patients undergoing tumour resection surgery at the Department of Neurosurgery (Erasmus University Medical Center, Rotterdam, The Netherlands). A written informed consent was signed by all patients accordingly to the Helsinki Declaration.

− Patient 1 was a 52-year-old-male with a tumour in the right temporal lobe secondary to a melanoma. There were no episodes of epilepsy or mental illness associated. The patient did not receive antiepileptic medication.
− Patient 2 was a 62-year-old male with a glioblastoma on the right temporal lobe. The patient presented epilepsy secondary to the tumour.
− Patient 3 was a 32-year-old male with and oligodendroglioma on the right fronto-temporal lobe. The patient received treatment for epileptic seizures.
− Patient 4 was a 60-year-old male who underwent surgery due to a tumour in the right frontal lobe derived from a lung carcinoma. There was no past psychiatric history or presence of epilepsy or seizures.
− Patient 5. Was a 69-year-old female who showed signs of glioblastoma in the right temporo-parietal lobe. There was no history of mental illness or presence of epilepsy.

### Viral labelling

Homozygous TSC::cre mice were used for cell-type specific labelling using a cre-dependent adeno-associated virus (AAV) expression. Mice were injected at 4 weeks of age with pENN.AAV.CamKII.HI.GFP-Cre.WPRE.SV40 (Addgene viral prep # 105551-AAV9) in the right hemisphere to silence *Tsc1* gene and pENN.AAV9.CamKII0.4.eGFP.WPRE.rBG (Addgene viral prep # 105541-AAV9) in the left hemisphere for control conditions. Viruses were diluted to 1/6 and 1/10 respectively to obtain a titer of 1×10^12^ vg/mL, in phosphate-buffered saline (PBS) to obtained sparce and region-focused labelling.

Mice were anaesthetised using 5% isofluorante (O_2_ flow of 0.5 L/min) and maintained with 2% isofluorante and 0.55% oxygen during surgery. Mice were placed in a stereotaxic frame using a mouth bar (Soelting) for head fixation, and their body temperatures was maintained at 37°C during the whole procedure. Animals were subcutaneously injected with Temgesic (buprenorphine 0.5 mg/kg) and Xylocaine (100 mg/mL, AstraZeneca) was sprayed locally on the skull to reduce pain and inflammation. A scalp incision was made to access the skull and a craniotomy was performed at the following sites of injection (in mm): mPFC: +1.75 bregma, ±0.35 lateral, -1.9 dorsoventral. Viral injection (100 nL per injection site) was performed using a borosilicate glass micropipette controlled by a syringe pump (infusion speed 0.05 μL/min). At the end of the injection, the micropipette was sustained in place for 5 minutes before withdrawal. The incision was closed with skin-glue (Derma+flex). After surgery, mice recovered for 4-6 weeks before electrophysiological or immunohistochemical analysis were executed.

### Electrophysiology

#### Mice

Mice were anaesthetised using 5% isoflurane. After decapitation, brains were removed in ice-cold partial sucrose-based solution containing (in mM): sucrose 70, NaCl 70, NaHCO_3_ 25, KCl 2.5, NaH_2_PO_4_ 1.25, CaCl_2_ 1, MgSO_4_ 5, sodium ascorbate 1, sodium pyruvate 3 and D(+)-glucose 25 (carboxygenated with 5% CO_2_/95% O_2_). Coronal slices from the prefrontal and somatosensory cortex (300 µm thick) were obtained with a vibrating slicer (Microm HM 650V, Thermo Scientific) and incubated for 45 min at 34 °C in holding artificial cerebro-spicnal fluid (ACSF) containing (in mM): 127 NaCl, 25 NaHCO_3_, 25 D(+)-glucose, 2.5 KCl, 1.25 NaH_2_PO_4_, 1.5 MgSO_4_, 1.6 CaCl_2_, 3 sodium pyruvate, 1 sodium ascorbate and 1 MgCl_2_ (carboxygenated with 5% CO_2_/95% O_2_). Next, the slices recovered at room temperature for another 15 min.

Slices were then transferred into the recording chamber where they were continuously perfused with recording ACSF (in mM): 127 NaCl, 25 NaHCO_3_, 25 D-glucose, 2.5 KCl, 1.25 NaH_2_PO_4_, 1.5 MgSO_4_ and 1.6 CaCl_2_. Cells were visualized using an upright microscope (BX51WI, Olympus Nederland) equipped with oblique illumination optics (WI-OBCD; numerical aperture 0.8) and a 40x water-immersion objective. Images were collected by a CCD camera (CoolSMAP EZ, Photometrics) regulated by Prairie View Imaging software (Bruker). In WT mice, layer II-III pyramidal cells in the frontal and somatosensory cortex were distinguishable by their morphological traits. In TSC^hom^ mice, transfected neurons were visualized by eGFP expression using a GFP filter (Semrock, Rochester, NY, USA).

Electrophysiological recordings were acquired using HEKA EPC10 quattro amplifiers and Patchmaster software (10 Hz sampling rate) at 33°C. Patch pipettes were pulled from borosilicate glass (Warner instruments) with an open tip of 3.5-5 MΩ of resistance and filled with intracellular solution containing (in mM) 125 K-gluconate, 10 NaCl, 2 Mg-ATP, 0.2 EGTA, 0.3 Na-GTP, 10 HEPES and 10 K2-phosphocreatine, pH 7.4, adjusted with KOH (280 mOsmol/kg), with 5 mg/mL biocytin to fill the cells. Series resistance was kept under 20 MΩ with correct bridge balance and capacitance fully compensated; cells that exceeded this value were no included in the study. Cells were filled with biocytin for at least 20 min.

Data analysis was performed offline using with AxoGraph X Office software (v1.7.0, AxoGraph Scientific). Physiological characteristics were determined from voltage responses to current injection pulses of 500 ms duration in 100 pA intervals ranging from -100 to +600. Action potential (AP) characteristics were obtained by the first elicited AP in the voltage response; AP peak was defined as the maximum peak of the AP; AP half-width was measured as the half of the peak amplitude. AP threshold was determined as the first inflection point where the rising membrane potential exceeded 50mV/ms slope. AP rise time was quantified as duration from 10% to 90% of the peak amplitude. AP decay time was measured as the duration from 100% to 50% of the peak. The afterhyperpolarisation (AHP) amplitude was measured as the hyperpolarizing peak from the initiation of the refractory period until the recovery state. AP frequency was estimated by the inverse of the difference between two consecutive AP in each current step. Likewise, passive membrane properties such as input resistance or conductance were calculated by the slope of the linear regression through the voltage-current curve. EPSC were recorded at -70 mV holding potential during 5 min. Synaptic events were analysed using Mini Analysis Program (Synaptosoft, Decatur, GA).

#### Human

*Ex vivo* human recordings of acutely resected cortex slices were obtained by overlying tissue that was removed in order to gain access to the tumour. After resection, the tissue block was transferred into carboxygenated (95% O_2_/5% CO_2_) ice-cold solution and sliced into 300 µm thick slices for electrophysiology. For the electrophysiological recordings, only slices where there was no infiltrating tumour found were utilized. Whole-cell recordings and data analysis were performed and analysed identically as the mouse tissue, except for current injection pulses that range from -100 to +600 in 50 pA steps.

### Immunohistochemistry

#### Mice slices

Mice were anaesthetized intraperitoneally with pentobarbital natrium. Midline skin incision was made at the thoracic outlet followed by opening of the abdomen and a cut in the diaphragm and the costal cartilage for cardial perfusion. The needle was placed into the apex of the left ventricle and animal was perfused with 4 % formaldehyde (PFA). Brains were dissected and fixed in 4% PFA for 2 h at room temperature. Next, brains were transferred into 10% sucrose phosphate buffer (0.1 M PB) and stored overnight at 4°C. For better handling and slicing of the brains, they were embedded in 12% gelatin-10% sucrose blocks and left during 1.5 h at room temperature in 10% paraformaldehyde-30% sucrose solution (PB 0.1 M) and later incubated overnight in 30% sucrose solution at 4°C.

Coronal sections (40 µm thick) were sliced using a freezing microtome (Leica, Wetzlar, Germany; SM 2000R) and stored in 0.1 M PB. Sections were blocked with 0.5% Triton X-100% (MerkMillipore) and 10% normal horse serum (NHS; Invitrogen, Bleiswijk, The Netherlands) for 1 h at room temperature, and incubated over 72 h at 4°C with mouse anti-NeuN (1:300, Millipore, MAB377), rabbit anti-pS6^S235/236^ (1:300, Cell Signaling Technologies, 2211S) in PBS buffer containing 0.4% Triton X-100 and 2% NHS. NeuN was visualized using anti-mouse Alexa647 secondary antibody (1:300, Invitrogen); pS6 was incubated in secondary antibody anti-rabbit Cy3 (1:300, Invitrogen). Secondary antibodies were incubated at room temperature for 2 h in PBS buffer containing 0.4% Triton X-100 and 2% NHS. Next, sections were washed with PBS and coverslipped in Vectashield H1000 fluorescent mounting medium (Vector Labs, Peterborough, UK).

#### Mice biotin-filled cells

Layer II-III pyramidal neurons were filled with 5mg/mL biocytin during whole-cell recordings and then fixed with 4% (PFA) overnight and stored in PBS at 4°C. Slices were rinsed with PBS and then stained with streptavidin-Cy3 secondary antibody (1:300; Invitrogen), 0.4% Triton X-100 and 2% NHS in PBS during 3 h. Slices were mounted on slides and coverslipped with 150 µl Mowiol (Sigma). After imaging with confocal at 63x, cells were unsealed and rinsed with PBS. To prevent thinning and dehydration, the slices were left in 30% sucrose overnight before resectioning. Using a freezing microtome, sections were recut into 40 µm slices and stored at 0.1 M PB. Resectioned slices were incubated using primary mouse anti-MBP (1:300) and secondary anti-mouse Alexa-647 (1:300) for eGFP cells and anti-mouse Alexa-488 (1:300) for WT mice.

#### Human

Human pyramidal cells were filled with 5mg/mL biocytin and fixed in 4% PFA overnight. Slices were washed with PBS and stained with streptavidin-Cy3 secondary antibody (1:300) in a PBS-based solution containing 0.4% Triton X-100 and 5% bovine serum albumin (BSA; Sigma-Aldrich, The Netherlands) during 3 h at room temperature. Slices were mounted and coverslipped. Slices were recut in 40 µm slices and then blocked in PBS containing 0.4% Triton X-100 and 5% BSA and posteriorly stained using mouse anti-MBP (1:300) in PBS, 0.4% Triton X-100 and 5% BSA during 72 h at 4°C. Secondary antibodies Alexa-488 anti-mouse and streptavidin-Cy3 were diluted in PBS, 0.4% Triton X-100 and 5% BSA during 3 h at room temperature. Next, slices were coverslipped and sealed for imaging.

### Confocal imaging and reconstruction

Images were taken using a Zeiss LSM 700 microscope (Carl Zeiss) equipped with Plan-Apochromat 10x/0.45 NA, 40x/1.3 NA (oil immersion) and 63x/1.4 NA (oil immersion) objectives. DAPI, Alexa488, Cy3 and Alexa647-secondary fluorophores were imaged using excitation wavelengths of 405 nm, 488 nm, 555 nm, and 639 nm, respectively.

In WT and human cells, a 10x magnification picture of the cell was taken to know exact distance from soma to pia, and a whole-cell overview image was acquired using 63x magnification objective and 555 nm wavelength. Pinhole was kept at 0.2% and gain was set at 750-800 units to adjust the signal to noise ratio. Biocytin-filled cells were imaged with tiled *z*-stack images (512 × 512 pixels) with a step size of 1 μm. Resectioned slices of such cells were obtained at 40x magnification using 488 and 555 nm wavelength filters.

For TSC^f/f^ eGFP^+^ cells a similar approach was used. A 10x magnification picture was taken to know location of the cell. To classify the cells in TSC1-specific deletion or control eGFP, a 63x magnification picture with *z*-stack (1 μm step size) and filters 555 nm and 488 nm was taken. Axonal examination of virally-labelled cells was obtained by 63x magnification tiled *z*-stack images (512 × 512 pixels) with a step size of 1 μm using 555 nm wavelength. MBP staining pictures were acquired using 40x magnification (1024 x 1024 pixels) and excitation wavelengths of 647 and 555 nm. The same settings were maintained across all pictures to ensure fluorescence was equally measured.

Overview images were then transferred into Neurolucida 360 software (v2.8; MBF Bioscience) and the axon was reconstructed using the interactive tracing with the Directional Kernels method. Reconstruction of the axon and myelinated segments were analysed with Neurolucida Explorer (MBF Bioscience). Distance to the pia and internodes were quantified and observed using ImageJ (ImageJ 5.12h). Axons were accepted as myelinated when they presented at least one MBP-positive internode and unmyelinated when no MBP-positive internode could be identified up to at least 10^th^ branch order. The distance to the first branch point was calculated as the length of the axon from the soma to the first branching axon segment. The distance to the first myelinated segment was determined as the distance along the axon from the soma to the initial point of the MBP segment.

For quantification of pyramidal-specific deletion of *Tsc1* as well as cell size analyses, one-tiled z-stack confocal picture was taken with 40x magnification. Stacks were randomly sampled across layer II-III of the targeted region. Three stacks were quantified per mouse and hemisphere. Only cells were pS6^S235/236^ was overlapping eGFP in the soma were used. Total immunofluorescent was determined pS6^S235/236^ as corrected total cell fluorescence (CTCF), with the following equation CTCF = integrated density – (area of selected cell x mean fluorescence of background readings). Randomly selected cells were manually outlined, and area and integrated density were obtained using ImageJ. To assess the exact area of the cell, NeuN marker was used in ImageJ to outline the cell and measure its area. No spatial corrections were made for tissue shrinkage.

### Structured illumination microscopy (SIM)

Structured illumination imaging was performed using a Zeiss Elyra PS1 system (Carl Zeiss). 3D-SIM data was acquired using a 63x/1.4 NA oil immersion objective. A 561 nm 100 mW diode laser together with a BP 570–650 + LP 750 filter was used to excite the fluorophores. Five phases and five rotations modulated the grating that was present in the light path during 3D-SIM acquisition. Serial z-stacks of 110 nm were recorded on an Andor iXon DU 885, 1002 × 1004 EMCCD camera. Raw data was reconstructed using Zen 2012 software (Zeiss) and posteriorly analysed in Fiji image analysis software.

In order to avoid overexposure from the soma, the first picture was taken starting from the second branch order axonal segment and continuing along this one. Axonal segments were imaged from branch point to the subsequent branch point. Bandwidth, offset and exposure time were regulated manually per segment to prevent oversaturation of the picture. Reconstructed pictures were loaded into Fiji and analysed using a custom-made script as previously described by *Stedehouder, J. et al* ^19^.

### Statistical analysis

All statistical analysis were operated using GraphPad 8.01. Data was firstly analysed for normality using Kolmogorov–Smirnov test. No outlier was identified or removed. Data sets following normal distribution were analysed using unpaired two-tailed *t*-test. Data sets without a normal distribution were analysed using Mann Whitney test.

A self-custom-made algorithm for R was created to calculate the receiver operating characteristic (ROC) curve, the area under the curve (AUC), the thresholds from the pure length and diameter univariate models and for the bivariate model. The optimum thresholds were determined as the maximum point of the Youden’s J statistic, calculated as the sum of the sensitivity and specificity minus 1. The AUC were estimated as the integral of the ROC curves for the univariate and bivariate models.

## Supplementary data

**Supplementary figure 1.**
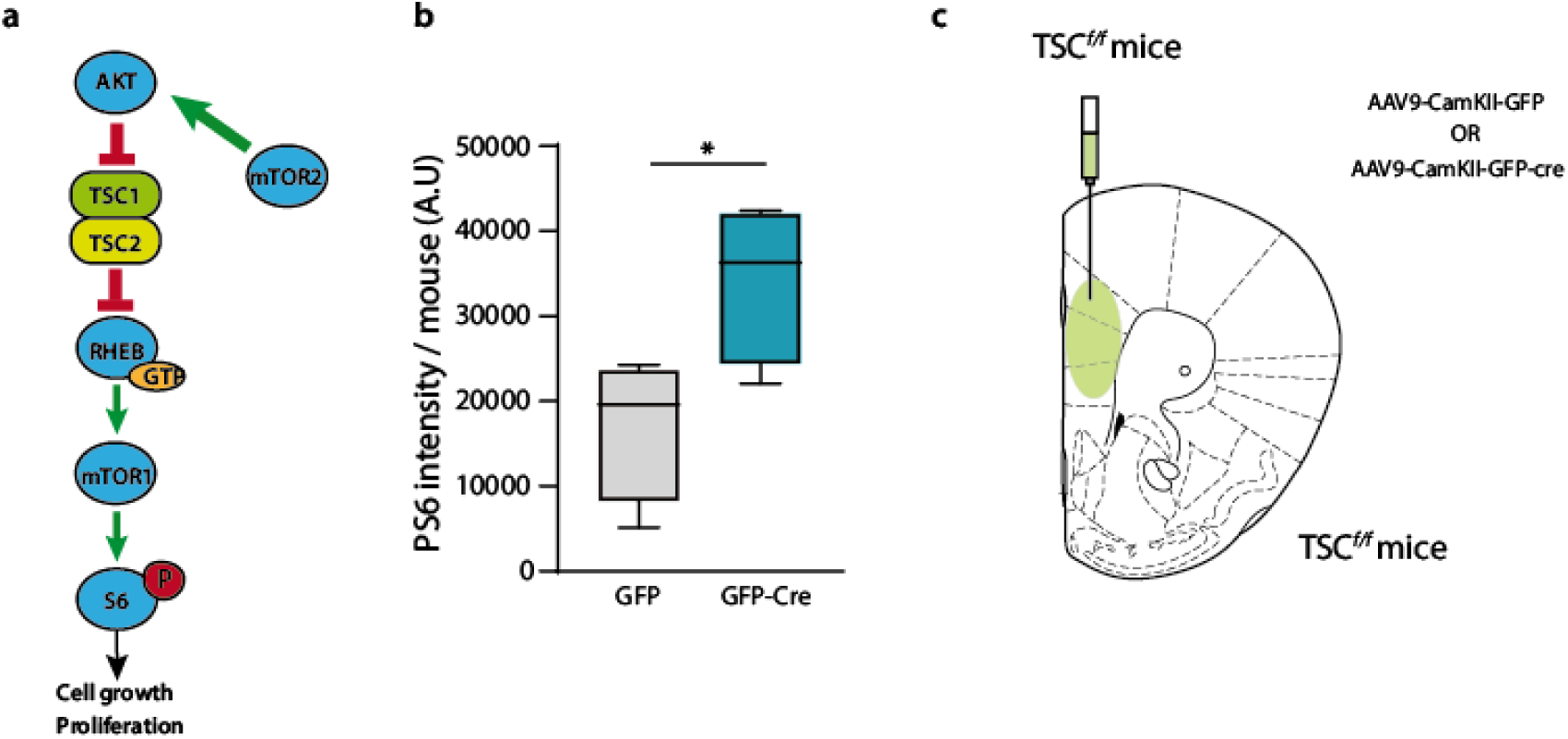
TSC1 deletion activates the mTOR pathway. **a)** mTOR signalling pathway. Green arrows indicate activation and red lines indicate inhibition. When TSC1 is knocked-out the TSC complex cannot be formed and RHEB binds a GTP molecule. In this manner, mTOR becomes active leading to the phosphorylation of ribosomal protein S6 that enables the transduction of growth and proliferation genes. **b)** Box plot of the area of the soma of PS6- and GFP-positive cells (t-test: t = 2.68; p-value = 0.036; GFP average: 17161 AU, n = 163 cells from 4 mice; GFP-Cre average: 34247 AU, n = 207 cells from 4 mice). **c)** Drawing of the viral injections in the S1 of TSC^f/f^ mice.

**Supplementary figure 2a.**
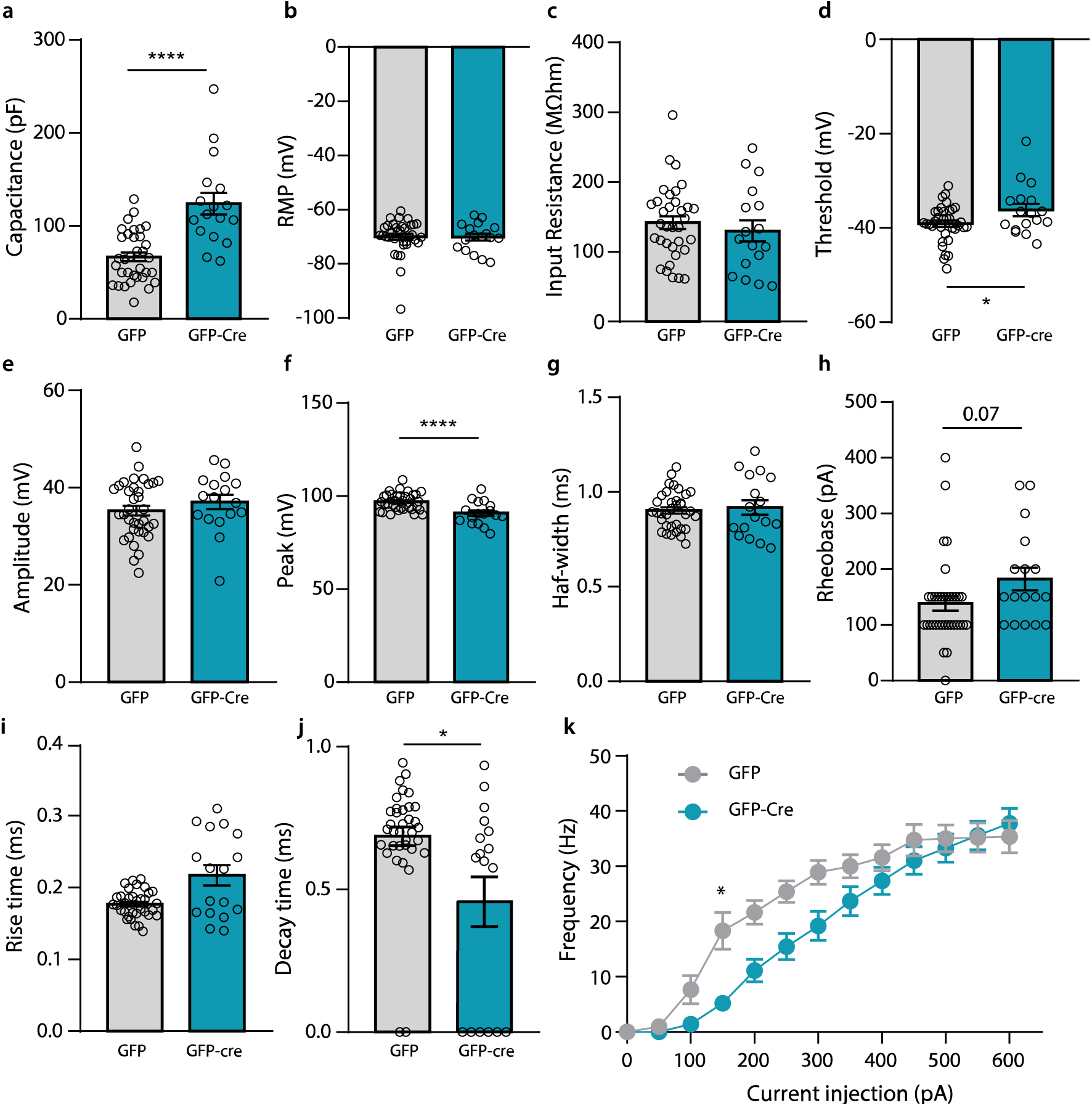
Electrophysiological properties of Cre lines in PFC. **a)** Capacitance of GFP-Cre cells was increased (t-test: 5.47; p-value < 0.0001; GFP average: 66.68 pF; GFP-Cre average: 123.8 pF). **b)** AP threshold was reduced (t-test: t = 2.24; p-value = 0.029; GFP average: -39.09 mV; GFP-Cre average: -36.17 mV). **c)** No variations in the RMP (t-test: t = 0.07; p-value = 0.94; GFP average: -70.02 mV; GFP-Cre average: -70.15 mV). **d)** Input resistance was unchanged (t-test: t = 0.71; p-value = 0.48; GFP average: 141.8 MΩ; GFP-Cre average: 129.9 MΩ). **e)** Amplitude was not changed (t-test: t = 0.99; p-value = 0.33; GFP average: 35.3 mV; GFP-Cre average: 37.04 pA). **f)** Peak was reduced in GFP-Cre compared to their control littermates (t-test: t = 4.034; p-value = 0.0002; GFP average: 96.91 mV; GFP-Cre average: 90.83 mV. **g)** Half-width was similar between groups (t-test: t = 0.422; p-value = 0.67; GFP average: 0.90 ms; GFP-Cre average: 0.92 ms. **h)** There was a trend towards higher rheobase (t-test: t = 1.86; p-value = 0.068; GFP average: 138.6 pA; GFP-Cre average: 182.4 pA). **i)** Rise time was increased (Mann-Whitney: U = 200; p-value = 0.058; GFP average: 0.18 ms; GFP-Cre average: 0.23 ms). **j)** Decay time was reduced (Mann-Whitney: U = 177; p-value = 0.018; GFP average: 0.71 ms; GFP-Cre average: 0.61 ms). **k)** Firing frequency in response to hyperpolarising and depolarising current steps.

**Supplementary figure 2b.**
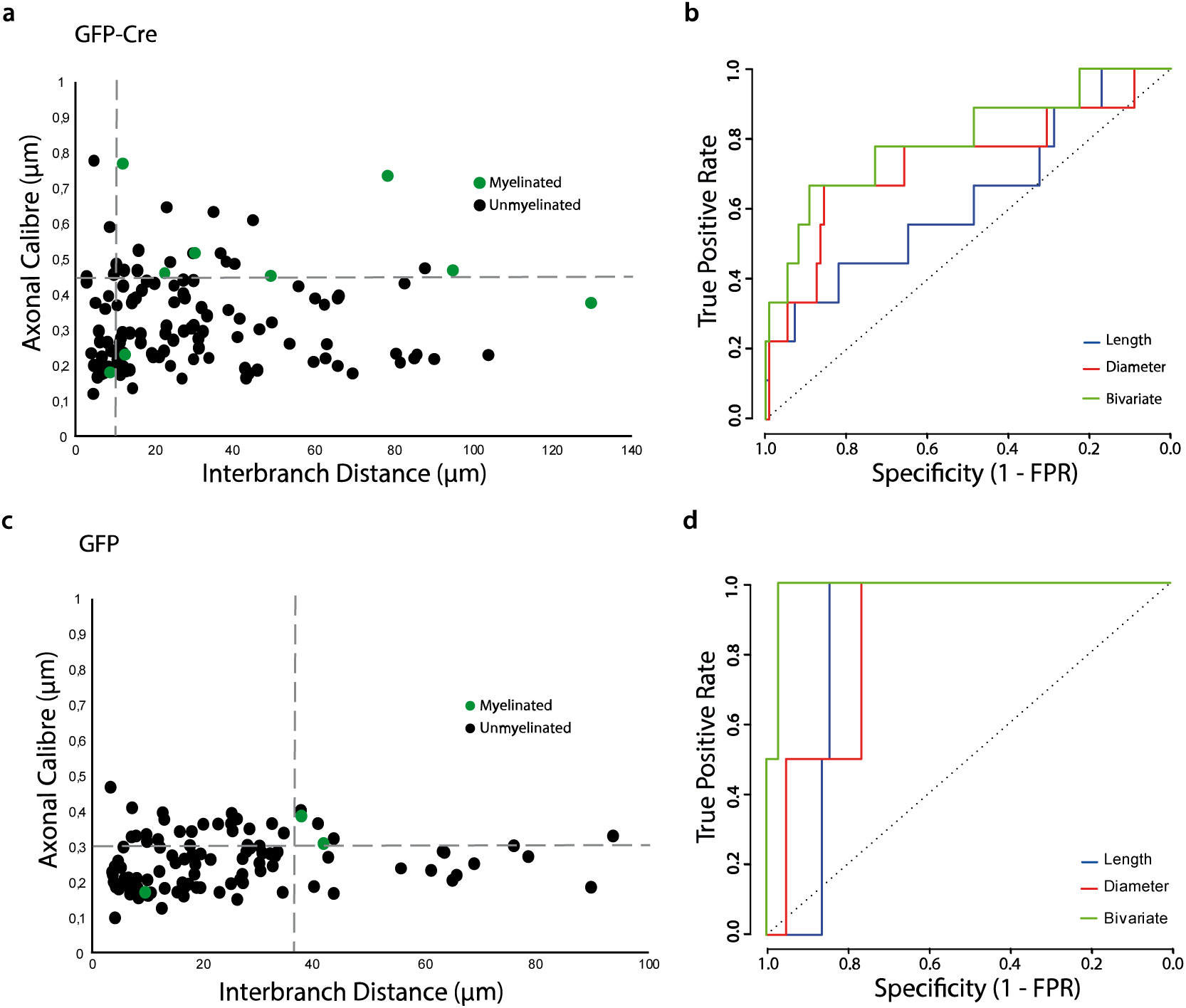
Axonal thresholds of Cre lines in PFC. **a)** Distribution of the axonal segment calibre and interbranch distances for myelinated (green) and unmyelinated (black) segments in GFP-Cre cells. Dotted lines depict the critical thresholds of the bivariate analysis. **b)** ROC curves for univariate length (length threshold = 47.55; AUC = 0.63; sensitivity = 0.44; specificity = 0.82), diameter (diameter threshold = 0.45; AUC = 0.73; sensitivity = 0.67; specificity = 0.86) and bivariate (joint threshold: length = 10.242, diameter = 0.45; AUC = 0.80; sensitivity = 0.67; specificity = 0.90) analyses of GFP-Cre axonal segments. **c)** Distribution of the axonal segment calibre and interbranch distances for myelinated (green) and unmyelinated (black) segments in GFP cells. Dotted lines depict the critical thresholds of the bivariate analysis. **d)** ROC curves for univariate length (length threshold = 37.21; AUC = 0.84; sensitivity = 1; specificity = 0.84), diameter (diameter threshold = 0.31; AUC = 0.86; sensitivity = 1; specificity = 0.76) and bivariate (joint threshold: length = 37.25, diameter = 0.30; AUC = 0.99; sensitivity = 1; specificity = 0.97) analyses of GFP axonal segments.

**Supplementary table 1.** Electrophysiological properties of PFC layer II-III pyramidal cells in WT mice at 8-12 weeks of age.

|  | MEAN | SEM |
| --- | --- | --- |
| Capacitance (pF) | 62,19 | 8,32 |
| RMP (mV) | -72,63 | 1,46 |
| Rinput (MΩ) | 202,78 | 22,04 |
| AP threshold (mV) | -37,93 | 0,71 |
| AP amplitude (mV) | 32 | 1,84 |
| AP peak (pA) | 92,31 | 2,01 |
| AP half-width (ms) | 0,99 | 0,08 |
| AP rise time (ms) | 0,20 | 0,02 |
| AP decay time (ms) | 0,76 | 0,07 |
| AP threshold (at 300 pA) (Hz) | 29,77 | 2,29 |
| Rheobase (pA) | 123,68 | 17,67 |

**Supplementary table 2.**
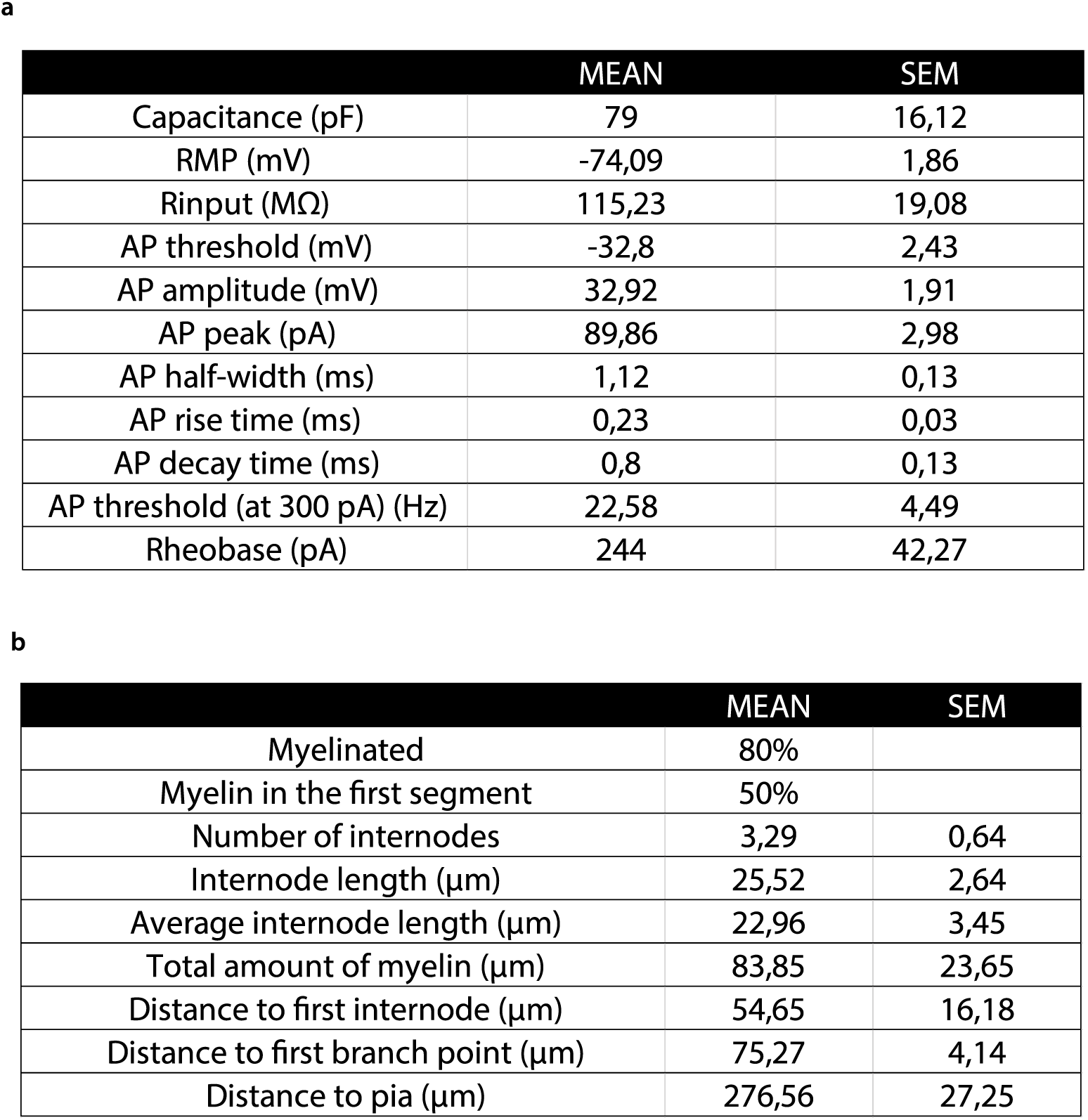
Electrophysiological properties and myelin profile of S1 layer II-III pyramidal cells in mice. **a)** Intrinsic properties of S1 pyramidal cells. **b)** Myelin characteristics of S1 pyramidal cells.

**a**
|  | MEAN | SEM |
| --- | --- | --- |
| Capacitance (pF) | 79 | 16,12 |
| RMP (mV) | -74,09 | 1,86 |
| Rinput (MΩ) | 115,23 | 19,08 |
| AP threshold (mV) | -32,8 | 2,43 |
| AP amplitude (mV) | 32,92 | 1,91 |
| AP peak (pA) | 89,86 | 2,98 |
| AP half-width (ms) | 1,12 | 0,13 |
| AP rise time (ms) | 0,23 | 0,03 |
| AP decay time (ms) | 0,8 | 0,13 |
| AP threshold (at 300 pA) (Hz) | 22,58 | 4,49 |
| Rheobase (pA) | 244 | 42,27 |

|  | MEAN | SEM |
| --- | --- | --- |
| Myelinated | 80% |  |
| Myelin in the first segment | 50% |  |
| Number of internodes | 3,29 | 0,64 |
| Internode length (μm) | 25,52 | 2,64 |
| Average internode length (μm) | 22,96 | 3,45 |
| Total amount of myelin (μm) | 83,85 | 23,65 |
| Distance to first internode (μm) | 54,65 | 16,18 |
| Distance to first branch point (μm) | 75,27 | 4,14 |
| Distance to pia (μm) | 276,56 | 27,25 |

**Supplementary table 3.** Electrophysiological and myelination properties of human pyramidal cells. **a)** Intrinsic properties of human pyramidal cells. **b)** Myelin characteristics of the cells that were myelinated. Correlation between electrophysiology and myelination pattern could not be done because of the low n of unmyelinated cells.

|  | MEAN | SEM |
| --- | --- | --- |
| Capacitance (pF) | 78,99 | 19,91 |
| RMP (mV) | -68,53 | 2,11 |
| Rinput (MΩ) | 189,1 | 34,67 |
| AP threshold (mV) | -40,79 | 3,25 |
| AP amplitude (mV) | 41,38 | 2,32 |
| AP peak (pA) | 93,99 | 3,14 |
| AP half-width (ms) | 0,96 | 0,03 |
| AP rise time (ms) | 0,24 | 0,01 |
| AP decay time (ms) | 0,67 | 0,10 |
| AP threshold (at 300 pA) (Hz) | 33,03 | 3,02 |
| Rheobase (pA) | 95 | 19,91 |

**Supplementary table 3. Electrophysiological and myelination properties of human pyramidal cells.**
|  | MEAN | SEM |
| --- | --- | --- |
| Myelinated | 89% |  |
| Myelin in the first segment | 38% |  |
| Number of internodes | 2,87 | 1,25 |
| Internode length (μm) | 49,37 | 12,67 |
| Average internode length (μm) | 41,31 | 17,68 |
| Total amount of myelin (μm) | 162,21 | 82,3 |
| Distance to first internode (μm) | 149,68 | 27,14 |
| Distance to first branch point (μm) | 137,36 | 62,73 |
| Distance to pia (μm) | 833,37 | 251,9 |

